# “Correcting” gene trees to be more like species trees increases topological error when incomplete lineage sorting is high

**DOI:** 10.1101/2022.08.21.504711

**Authors:** Zhi Yan, Huw A. Ogilvie, Luay Nakhleh

## Abstract

The evolutionary histories of individual loci in a genome can be estimated independently, but this approach is error-prone due to the limited amount of sequence data available for each gene, which has led to the development of a diverse array of gene tree error correction methods which reduce distance to the species tree. We investigate the performance of two representatives of these methods: TRACTION and TreeFix, in the case where incomplete lineage sorting is high. We found that gene tree error correction only increases the level of error in gene tree topologies by “correcting” them to be closer to the species tree, even when the true gene and species trees are discordant. We confirm that full Bayesian inference of the gene trees under the multispecies coalescent model is more accurate than independent inference. Gene tree correction must be considered a multi-locus task where the gene tree distribution is taken into account, rather than treating gene trees independently.

**Significance statement:** Gene tree information is essential for studying elucidating gene, genome, species, and phenotypic evolution, and a wide array of phylogenetic methods have been developed for gene tree estimation. Given that gene tree estimates are often inaccurate, several methods for “correcting” gene tree estimates have been devised. Here we show that correction methods that neglect the distribution of gene trees that is induced by the species phylogeny could produce poor results, calling for the development of species phylogeny-aware gene tree correction.

## Introduction

Although genes evolve within the context of species, the evolutionary history of genes and gene families are unique and different from the species phylogeny because of processes such as incomplete lineage sorting, gene duplication and loss (GDL), and horizontal gene transfer (HGT) (Maddison 1997). Deciphering these individual histories of particular gene families is of great interest; to pick a few discoveries enabled by inferred gene trees, they have revealed effector and resistance genes in plant-pathogen interactions (McDonald *et al*. 2016; Yang *et al*. 2013), supported the importance of visual system changes to the adaptive radiation of cichlids (Torres-Dowdall *et al*. 2015) and identified orthologs of genes linked to human health and disease in model organisms (Maxwell *et al*. 2014; Waaijers *et al*. 2015).

Now that sequencing and assembly of eukaryotic genomes is relatively routine (Michael and VanBuren 2020; Rhie *et al*. 2021), in aggregate an enormous amount of data is available for phylogenetic analyses. Using the megabases or gigabases (Oliver *et al*. 2007) available in each genome, precise and accurate species histories can be inferred (Hahn and Nakhleh 2016). However, to infer the history of individual gene families, the amount of information is much more limited with an average eukaryotic coding sequence length of roughly 1.3 kilobases (Xu *et al*. 2006). Fortunately, this limited sequence data can be augmented by information from the species phylogeny. When genes evolve following the multispecies coalescent (MSC) model, joint inference of the species and gene trees is substantially more accurate than inferring gene phylogenies independently (Szöllősi *et al*. 2014).

While joint inference methods are available for MSC (e.g. StarBEAST2; Ogilvie *et al*. 2017) or duplication and loss models (e.g. PHYLDOG; Boussau *et al*. 2013) of gene evolution, such methods are computationally intensive (Ogilvie *et al*. 2016). This has spurred the development of gene tree error correction tools intended to deal with GDL and HGT (David and Alm 2011; Jacox *et al*. 2016; Nguyen *et al*. 2012; Schreiber *et al*. 2014; Durand *et al*. 2005; Lai *et al*. 2017; Sjöstrand *et al*. 2012, 2014; Noutahi *et al*. 2016; Rasmussen and Kellis 2010; Wu *et al*. 2013; Bansal *et al*. 2018; Morel *et al*. 2020), which were developed to improve independently inferred gene trees through the reconciliation with a given species tree. These approaches are more scalable and trivially parallelizable.

As has been appreciated for decades, in shallow phylogenies, ancestral polymorphism can persist through speciation events, leading to incomplete lineage sorting (ILS) which is one of the major sources of gene tree heterogeneity (Suh *et al*. 2015; Wang *et al*. 2018; Alda *et al*. 2019). It is even possible that there are regions where the most probable gene tree topology differs from the species tree (Degnan and Rosenberg 2006). Failure to account for common outcomes of evolutionary processes, like ILS as an outcome of population genetics, is likely to yield misinterpretations of evolutionary history. Note that although there are existing gene tree error correction approaches allowing for ILS, they are either parsimony-based or non-parametric, not incorporating coalescent process probabilistically (Stolzer *et al*. 2012; Christensen *et al*. 2019, 2020).

In this study, we picked TreeFix (Wu *et al*. 2013) and TRACTION (Christensen *et al*. 2019, 2020) as two representative methods for species tree attraction based methods of gene tree error correction, hereafter species tree attraction methods. The former is a popular method utilizing the information from the species tree and sequence data based on a GDL model. The latter is a very recent non-parametric method that improves the uncertain branches by solving the RF-Optimal Tree Refinement problem, and it has been shown to be accurate when applied on simulated data with ILS. Our results show these methods performed badly when levels of ILS are high, due to a simplistic approach to error correction that reduces the distance between gene and species trees rather than considering the whole distribution of gene trees.

## Results

To understand how these species tree attraction methods would work on varying levels of GTEE, we simulated multiple sequence alignments with a broad range of informativeness by varying the number of sites and also studying a 10-fold change to the population mutation rate. These simulation settings resulted in six levels of GTEE spaced at roughly even intervals according to their average RF distances (Table 1).

**Table 1:** Levels of gene tree estimation error (GTEE) and trends

| $\theta$ | 0.001 | | | 0.01 | | |
| --- | --- | --- | --- | --- | --- | --- |
| #Sites | 200 | 600 | 1000 | 200 | 600 | 1000 |
| Avg. #Parsimony-informative sites | 0.17 | 0.51 | 0.86 | 1.78 | 5.11 | 8.58 |
| GTEE (Avg. RF) <sup>a</sup> | 0.689 | 0.537 | 0.453 | 0.324 | 0.201 | 0.148 |
| $[RF(TRACTION, ST) < RF(GT, ST)]$ % <sup>b</sup> | 89.4% | 76.3% | 63.4% | 47.0% | 27.9% | 20.0% |
| $[RF(TRACTION, ST) < RF(\widehat{GT}, ST)]$ % <sup>c</sup> | 94.1% | 79.8% | 67.0% | 49.1% | 26.6% | 18.5% |
| $[RF(TreeFix, ST) < RF(GT, ST)]$ % <sup>d</sup> | 89.1% | 82.1% | 82.3% | 79.9% | 73.1% | 60.8% |
| $[RF(TreeFix, ST) < RF(\widehat{GT}, ST)]$ % <sup>e</sup> | 94.9% | 85.7% | 85.5% | 81.3% | 73.1% | 60.5% |

<sup>a</sup> Gene tree estimation error, measured by the average RF distance between the true and IQ-TREE-inferred gene trees.
<sup>b</sup> Percentage of TRACTION-corrected gene trees that are closer in distance to the species tree than corresponding true gene trees.
<sup>c</sup> Percentage of TRACTION-corrected gene trees that are closer in distance to the species tree than corresponding IQ-TREE-inferred gene trees.
<sup>d</sup> Percentage of TreeFix-corrected gene trees that are closer in distance to the species tree than corresponding true gene trees.
<sup>e</sup> Percentage of TreeFix-corrected gene trees that are closer in distance to the species tree than corresponding IQ-TREE-inferred gene trees.

As our model species tree has 4 ingroup taxa, the methods could get 0, 1 or 2 splits correct, thus the distribution of unrooted RF distances has exactly three possible categories, and any shift in the distribution of distances to the right indicates an increase in GTEE, and conversely a shift to the left indicates a decrease (Fig. 1). Although TreeFix and TRACTION are both designed to improve the accuracy of estimated gene trees, our analysis shows that both methods actually *increased* GTEE under our simulation settings (Fig. 1). TreeFix performed worse than TRACTION; while TreeFix has been demonstrated good performance on data under GDL-only scenarios, the model misspecification of ignoring ILS clearly severely hurts the performance of that method (Fig. 1, Table 2).

**Figure 1:**
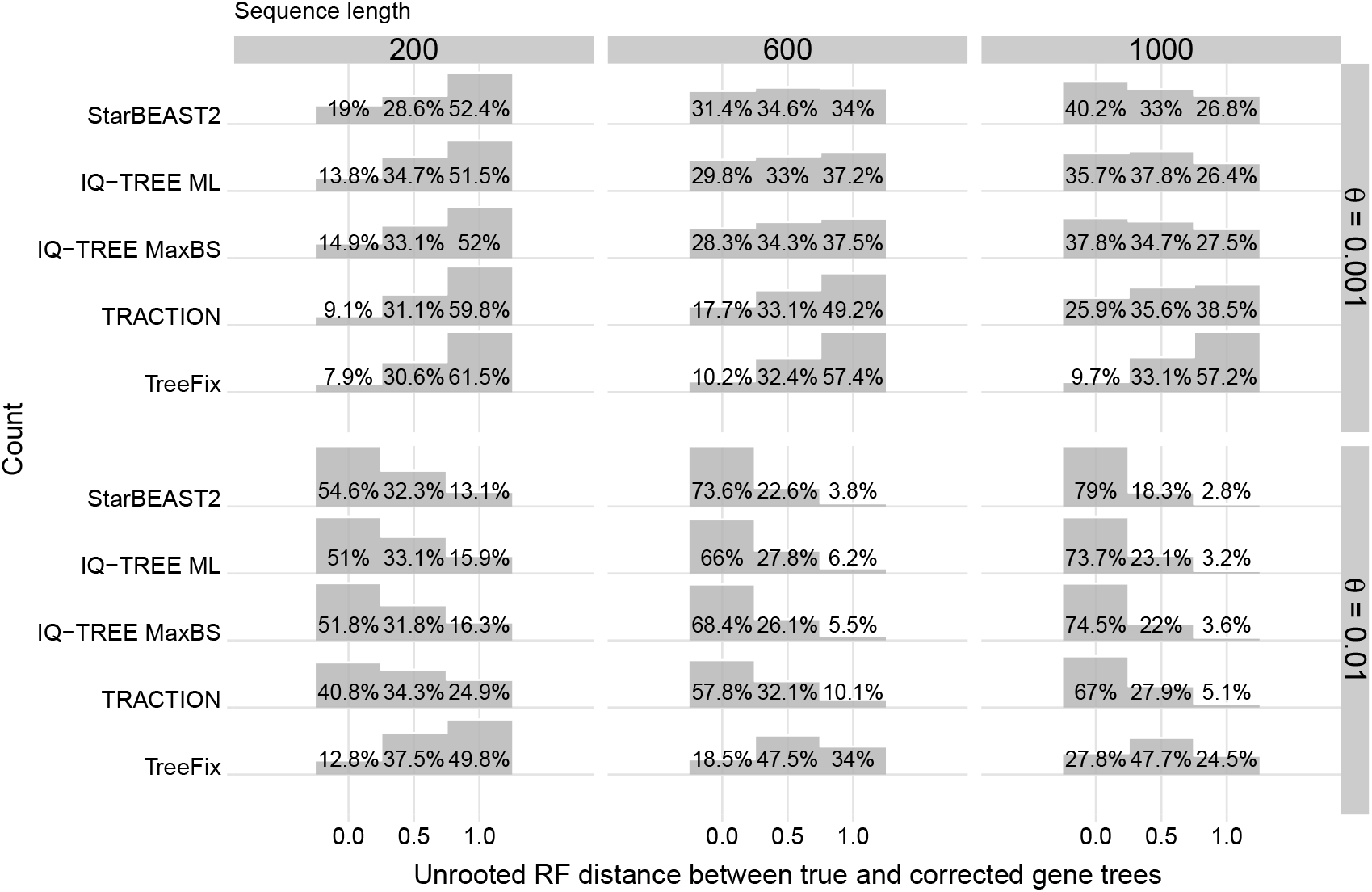
Error distributions of StarBEAST2-inferred, uncorrected and corrected gene trees. Uncorrected gene trees were either the maximum likelihood (ML) or maximum bootstrap support (MaxBS) topologies inferred using IQ-TREE. ML gene trees were corrected using either TRACTION or TreeFix. Gene trees from every replicate for all datasets simulated regardless of the number of loci were aggregated for each combination of population mutation rate *θ* and sequence length.

**Table 2:** Kullback-Leibler divergence between the theoretical distribution of gene tree topologies and either simulated or estimated distributions.

| $\theta$ | #Sites | TrueGT<sup>a</sup> | StarBEAST2 | IQ-TREE | MaxBS<sup>a</sup> | TRACTION | TreeFix |
| --- | --- | --- | --- | --- | --- | --- | --- |
| 0.001 | 200 | 0.0074 | 0.6419 | 1.1112 | 1.8330 | 3.5451 | 3.5348 |
| 0.001 | 600 | 0.0074 | 0.1994 | 0.4926 | 0.9673 | 2.4529 | 3.0026 |
| 0.001 | 1000 | 0.0074 | 0.0729 | 0.2374 | 0.4909 | 1.6643 | 2.9268 |
| 0.010 | 200 | 0.0074 | 0.0270 | 0.0434 | 0.1064 | 0.9101 | 2.4904 |
| 0.010 | 600 | 0.0074 | 0.0176 | 0.0109 | 0.0199 | 0.3568 | 1.8684 |
| 0.010 | 1000 | 0.0074 | 0.0099 | 0.0085 | 0.0092 | 0.1797 | 1.3363 |

<sup>a</sup> The distribution of simulated true gene trees, and that of the highest bootstrap-support trees.

We next examined the degree to which the TRACTION and TreeFix corrected gene trees towards the reference species tree, away from the true gene trees simulating using the MSC, or gene trees inferred from sequences simulated along those true gene trees. For both methods, the output topologies always had at least identical distances to the species tree than the uncorrected input topologies. In all cases except when TRACTION was applied to the higher population mutation rate gene trees, which had lower GTEE, the distances of most gene trees to the species tree were strictly smaller after correction (Table 1). Since 93.4% of the simulated genealogies here were in fact discordant with the species tree, this presumably only increased the GTEE.

The only approaches we found that decreased GTEE were bootstrapping or full Bayesian inference; the MaxBS tree consistently performed slightly better than the ML tree produced by IQ-TREE and gene trees inferred under the MSC using StarBEAST2 were substantially more accurate (Fig. 1). To make the latter case a fair comparison with TRACTION and TreeFix, the same true species tree was used to infer gene trees instead of joint inference.

## Discussion

The proliferation of gene tree error correction methods demonstrates the intense level of interest in the problem of improving gene tree accuracy without resorting to joint inference of species and gene phylogenies. However, we have demonstrated here that species tree attraction methods should be used with extreme caution when ILS causes the true gene tree histories to be highly discordant from the history of the corresponding species.

A previous investigation into gene tree error correction found that species tree attraction methods work well when uncorrected GTEE is high and gene tree discordance is much lower than in our analysis (Christensen *et al*. 2019, 2020). This is compatible with our finding that these methods essentially modify gene trees to be closer in distance to the species tree, since when ILS is relatively low the true gene trees will be more congruent with the species tree. So when GTEE is high the uncorrected gene trees will be somewhat random and simply altering them to be less random and more similar to the species tree will therefore improve accuracy. However, as we have shown, when ILS is higher, species tree attraction methods will increase GTEE regardless of the level of error in the originally inferred gene trees.

The only approaches we have found that decrease GTEE are to use the topology with the maximum bootstrap support, or full Bayesian inference under the MSC. The former approach is in a heuristic way marginalizing over branch lengths to derive the most likely topology, but is only minimally more accurate than simple ML inference. The latter approach results in substantially greater accuracy, as has been previously reported (Szöllősi *et al*. 2014). Given the poor scalability of full Bayesian MSC inference, and lack of ability to parallelize such inference, we believe it is still worth pursuing gene tree error correction methods. However existing methods are inadequate solutions when ILS is high, and future methods need to reflect the *distribution* of gene trees expected under a given model (whether the MSC or otherwise), rather than only considering the distance from the species tree, which may be discordant with many or most gene trees. We suggest that fixing gene trees should be a “multilocus task” and not carried out on individual loci without regard to the overall distribution.

## Materials and Methods

We used the five-taxon comb (fully unbalanced) tree from Liu and Edwards (2009) as our model species tree, keeping the branch lengths of the four-taxon ingroup clade while extending the outgroup branch, so that only the ingroup species resides in the anomaly zone. Following the simulation study of Wang and Nakhleh (2018), we performed coalescent simulations to generate datasets with 2, 5, 10, 20 and 50 gene trees, each with 10 replicates and a single individual per species. For each gene tree, we employed Seq-Gen (Rambaut and Grassly 1997), and used two different population mutation rates (*θ* = 4*Nµ*): 0.001 and 0.01, to simulate sequence data of length 200, 600 and 1000 nucleotides under the Jukes-Cantor substitution model (Jukes *et al*. 1969). Scaling the gene trees in this manner only changes the substitution rate *µ* and not the population size component *N* of *θ*.

The alterations of mutation rate and locus length amount to varying level of gene tree estimation error. In total, 5 × 10 × 2 × 3 = 300 datasets were produced. We then utilized IQ-TREE version 2.1.3 (Minh *et al*. 2020) with 100 bootstrap replicates under the Jukes-Cantor model to reconstruct maximum likelihood (ML) gene trees from these simulated alignments, and rooted them using the known outgroup. Furthermore, with these bootstrap replicates for each dataset, we obtained the most frequent topology, referred to as MaxBS (Maximum Bootstrap Support) tree.

We applied two programs TreeFix v1.1.10 (Wu *et al*. 2013) and TRACTION v1.0 (Christensen *et al*. 2019, 2020) to error-correct gene trees inferred by IQ-TREE. TRACTION requires a threshold value for contracting low support branch, so we adopted a support threshold of 75% as used in Christensen *et al*. (2019). Additionally, we ran StarBEAST2 (Ogilvie *et al*. 2017), a method for joint Bayesian inference of species and gene trees with fixed reference species tree, to sample posterior distributions of gene phylogenies from simulated sequences. We used, as point estimates of the gene trees, the maximum clade credibility tree topologies from the posterior distributions (Heled and Bouckaert 2013).

We defined and quantified gene tree estimation error using the unrooted normalized Robinson-Foulds distance (Robinson and Foulds 1981) between inferred uncorrected or corrected gene trees and the true simulated gene trees.

## Data Availability

The data underlying this article are available in the GitHub Repository, at https://github.com/Moerz/Gene-Tree-Fixing.

